# Subtle alterations in neonatal neurodevelopment following early or late exposure to prenatal maternal immune activation

**DOI:** 10.1101/2021.08.12.456095

**Authors:** Elisa Guma, Emily Snook, Shoshana Spring, Jason P. Lerch, Brian J. Nieman, Gabriel A. Devenyi, M. Mallar Chakravarty

## Abstract

Prenatal exposure to maternal immune activation (MIA) is a risk factor for a variety of neurodevelopmental and psychiatric disorders. The timing of MIA-exposure has been shown to affect adolescent and adult offspring neurodevelopment, however, less is known about these effects in the neonatal period. To better understand the impact of MIA-exposure on neonatal brain development, we first assess neonate communicative abilities with the ultrasonic vocalization task, followed by high-resolution *ex vivo* magnetic resonance imaging (MRI) on the neonatal (postnatal day 8) brain. Early exposed offspring displayed decreased communicative ability, while brain anatomy appeared largely unaffected, apart from some subtle alterations. By integrating MRI and behavioural assays to investigate the effects of MIA-expsoure on neonatal neurodevelopment we show that offspring neuroanatomy and behaviour are only subtly affected by both early and late exposure. This suggests that the deficits often observed in later stages of life may be dormant, not yet developed in the neonatal period, or not as easily detectable using a cross-sectional approach.

## 1. Introduction

*In utero* exposure to maternal infection is an environmental risk factor for neurodevelopmental and psychiatric disorders in exposed offspring(1–3). Maternal response to infection involves an increase in circulating proinflammatory cytokines. Although adaptive in terms of maternal health, this response may have unwanted consequences by interfering with the regulatory roles of these immune molecules in the developing fetal brain (4–8). Importantly, both maternal immune responsiveness and neurodevelopmental processes in the fetus vary across gestation; thus, the timing of maternal immune activation (MIA)-exposure in gestation may influence the nature and severity of anatomical and behavioural disruptions in offspring (9). Identifying which gestational windows are most sensitive may be critical to our understanding of the effects of MIA-exposure. Previously, our group has shown that the gestational timing of MIA-exposure has a differential impact on offspring neuro- and behavioural development at different developmental stages. In the embryo brain, we observed striking brain volume increases following MIA-exposure late in gestation (gestational day [GD]-17), while exposure in early gestation (GD9) resulted in more focal volumetric decreases (10). In our investigations of brain development trajectories from adolescence to adulthood, we observed early MIA-exposure to induce greater neuroanatomical and behavioural alterations than either late exposure or saline exposure, particularly in adolescence and early adulthood (11). This raises the question of what processes are occurring throughout the period between the embryo and childhood; namely: the neonatal period. Importantly, we previously identified this time period as being under-studied and receiving less attention in the MIA neuroimaging literature (12).

Human neuroimaging evidence provides more motivation for studying the neonatal period in more detail. Recent neonatal imaging studies have found that *in utero* exposure to increased maternal inflammation (measured with either interleukin 6 or C-reactive protein) was associated with alterations to both functional and structural connectivity in the infant brain as well as to toddler executive function (13–15). Some evidence of altered neonatal development following MIA-exposure exists in the animal literature as well. Positron emission tomography studies of the neonatal rabbit brain report increased neuroinflammation, as measured by TSPO, in MIA-exposed kits at postnatal day (PND) 1, and sustained until PND17 (16). MRI studies of MIA-exposed rhesus monkeys have also reported accelerated brain growth driven by white matter expansion in the first two years of life (17). Even though some observations in early phases of life exist (17–20), further work is required to understand the neurodevelopmental sequelae in the neonatal period following MIA-exposure, as this is a plastic window of brain development important for putative intervention.

To examine this sensitive period we employ high-resolution *ex vivo* structural magnetic resonance imaging (MRI), a technique with comparable signal across species (21) to examine the effects of *in utero* exposure to early (GD 9) or late (GD 17) MIA with the viral mimetic, polyinosinic:polycytidylic acid (poly I:C), on the neonate (PND8) brain. We also assayed neonate communicative ability using the ultrasonic vocalization (USV) task to phenotype neonatal mice at PND8. Much to our surprise, limited alterations were observed in neonate neuroanatomy due to either GD 9 or 17 exposure, while subtle decreases in communication were observed in the GD 9 exposed offspring.

### 2.1 Materials and methods

#### 2.1.1 Animals, prenatal immune activation, and sample preparation

Pregnant dams were bred in our animal facility using timed mating procedures in female and male C57BL/6J mice (8-12 weeks old) (described in **Supplement 1.1**). Pregnant dams were randomly assigned to one of four treatment groups (**Figure 1** for experimental design): (1) poly I:C (P1530-25MG polyinosinic:polycytidylic acid sodium salt TLR ligand tested; Sigma Aldrich) (5mg/kg, intraperitoneally) at GD 9 (POL E; 6 dams), (2) 0.9% sterile NaCl solution at GD 9 (SAL E; 5 dams), (3) poly I:C at GD 17 (POL L; 6 dams), or (4) saline at GD 17 (SAL L; 4 dams).

**Figure 1.**
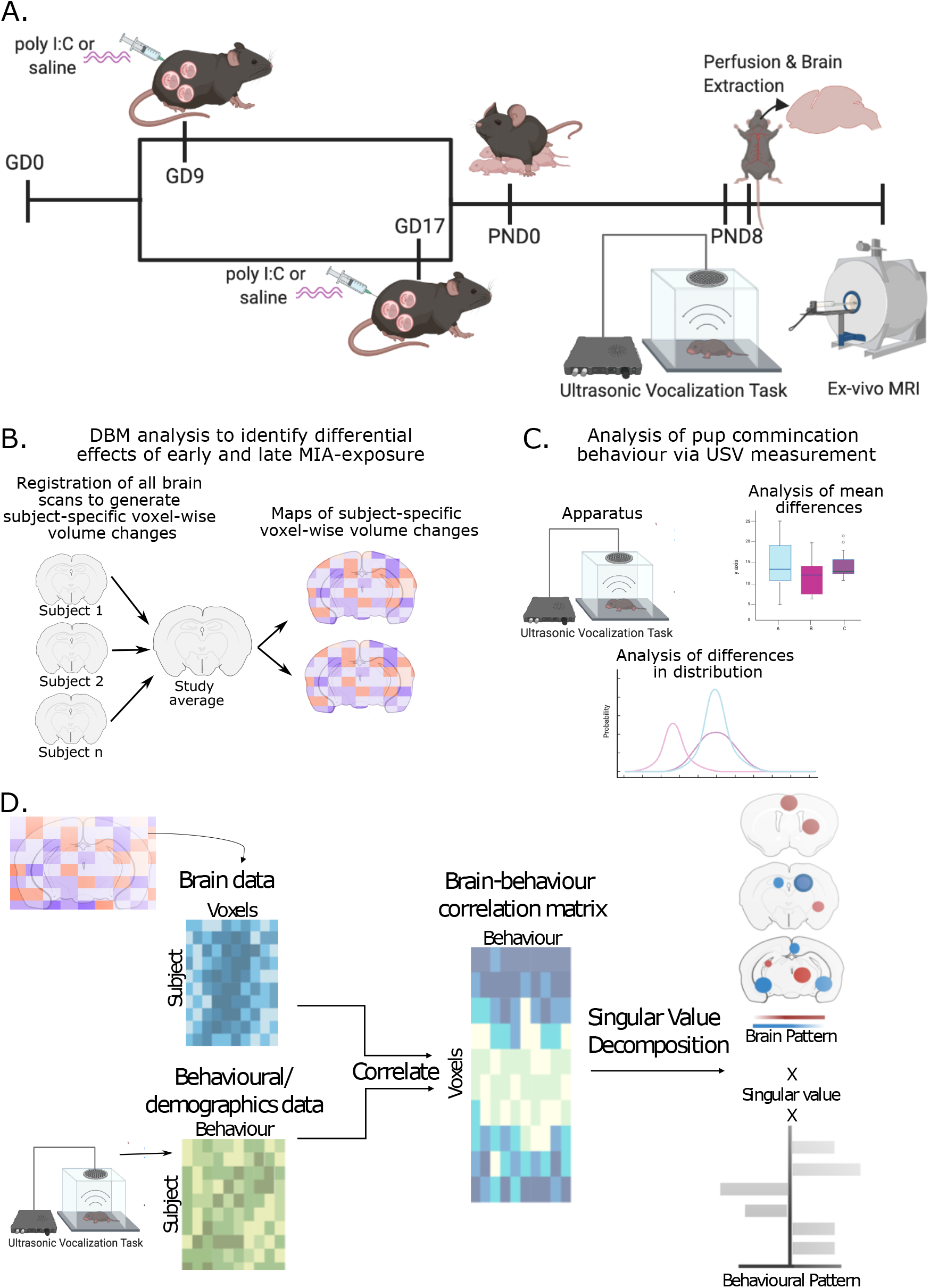
Experimental timeline. **A.** Pregnant dams were injected on either GD 9 or 17 with either poly I:C or vehicle (as in A). On PND8 offspring were tested on the ultrasonic vocalization task to assess communicative behaviour. Following behaviour, mice were perfused, and brains were prepared for *ex-vivo* MRI. **B.** Analysis flow of deformation-based morphometry analysis used to detect voxel-wise brain volume differences due to early or late MIA-exposure. **C.** Analysis of ultrasonic vocalization data for mean differences in call duration and frequency, and differences in distribution of calls. **D.** Multivariate analysis of brain-behavior data using PLS to identify patterns of covariation.

##### Neonatal Sample Preparation for MRI

On PND8, ~1 hour following behavioural testing (see **section 2.3**) neonates were perfused first with 20mL of 1X PBS, 2% gadolinium, and 1 μL/mL heparin (1000 USP units/mL) to remove any blood from the circulatory system, followed by 20mL of 4% PFA with 2% gadolinium in PBS solution. Skulls were dissected and stored in a PBS solution with 4% PFA and 2% gadolinium for 24 hours, after which they were transferred to a 0.02% Sodium Azide 1x PBS solution for long-term storage until scanning (see **Supplement 1.2** for more detail; sample numbers in **Table 1**). Immunostimulatory potential of poly I:C was confirmed in separate dams, all collected with the first batch of poly I:C (**Supplement 1.1, 2.1, Supplementary table 1**)(data from (10)).

### 2.2 Magnetic resonance image acquisition and processing

All samples were shipped to the Mouse Imaging Centre (Toronto, ON) for scanning. A multi-channel 7.0-T MRI scanner with a 40 cm diameter bore (Varian Inc., Palo Alto, CA) was used to acquire anatomical images of the neonate brains within skulls. A custom-built 16-coil solenoid array was used to acquire 40 μm^3^ resolution images from 16 samples concurrently (Dazai et al., 2011; Lerch et al., 2011) (see **Supplement 1.4** for details).

Preprocessed neonate brain images of all subjects in the study were aligned by unbiased deformation based morphometry using the antsMultivariateTemplateConstruction2.sh tool (https://github.com/CoBrALab/twolevel_ants_dbm)(22). The output of this iterative group-wise registration procedure is a study average against which groups can be compared, as well as deformation fields that map each individual subject to the average at the voxel level. Relative log-transformed Jacobian determinants (23), which explicitly model only the non-linear deformations and remove global linear transformation (attributable to differences in total brain size) were blurred at 160 μm full-width-at-half-maximum to better conform to Gaussian assumptions for downstream statistical testing.

### 2.3 Behavioural testing: ultrasonic vocalization task

Isolation-induced ultrasonic vocalizations of neonate offspring were assessed with standard procedures (24,25). This test was selected because, apart from motor abilities, it is one of the few complex behaviours that can be assessed this early in development (26,27) and because communicative abilities are a core deficit of ASD (28). Testing was performed using the Noldus UltraVox™ system (Noldus Information Technology, Leesburg, VA) on PND 8, as the rate of calling peaks around this time in mouse pups (Scattoni et al., 2008). Duration of each individual call per session per animal were recorded as raw data, as were the following summary measures per animal: total number of calls, maximum and minimum duration of calls, maximum and minimum call interval. For simplicity, call duration (and mean duration) was the only measure analyzed and discussed. See **Table 1** for sample size, and **Supplement 1.5** for details.

**Table 1.** Final sample size neonate MRI acquisition and neonate USV data following quality control.

| Group | Males (MRI) | Females (MRI) | Males (USV) | Females (USV) | Litters | # of pups per litter (median [range]) |
| --- | --- | --- | --- | --- | --- | --- |
| SAL E | 7 | 9 | 13 | 10 | 5 | 7 [2-8] |
| SAL L | 8 | 11 | 14 | 16 | 4 | 9 [5-9] |
| POL E | 13 | 13 | 23 | 17 | 6 | 7 [5-8] |
| POL L | 16 | 13 | 25 | 14 | 6 | 8 [3-10] |

### 2.4 Statistical analyses

#### 2.4.1 Neuroimaging data analysis

Statistical analyses were performed using the R software package (R version 3.5.1, RMINC version 1.5.2.2 www.r-project.org). First, we confirmed that there were no statistically significant differences between our two control groups (SAL E and SAL L), which allowed us to combine them into a single group, leaving us with three groups: saline (SAL), GD 9-poly I:C (POL E), and GD 17-poly I:C (POL L), consistent with our previous work (11). To assess the effects of poly I:C exposure either early or late in gestation on neonatal neuroanatomy, we ran a whole-brain voxel-wise linear mixed-effects model (lme4_1.1-21 package; (29)) on the relative Jacobian determinant files using group and sex as fixed effects, and number of pups per litter (ranging from 2-10; **Table 1**) as random intercepts. The litter size was selected as a random effect as it may have quite a significant effect on development in early phases of life (30); since we do not cull litters to be the same size, we considered this to be an important factor to control. The False Discovery Rate (FDR) correction was applied to correct for multiple testing (**Supplement 1.6.1** for details). This analysis was run again with the POL L group as the reference in order to directly compare POL E to POL L differences. Sex-by-group interactions were explored as a follow up analysis as described in **Supplement 1.6.1** and **Supplement 2.2.2**.

#### 2.5.2 USV data analysis

Since there were no differences in group means (see **Supplement 1.6.2** for analysis details), we used a hierarchical shift function in order to maximize the data collected for each mouse (31). This allows us to quantify how two distributions differ based on deciles of the distributions, i.e. it describes how each decile should be shifted to match one distribution to another. When a significant difference is observed between deciles it suggests that there is a specific difference in the number of calls between groups at a specific call length; this allows us to determine whether differences are consistent across the entire distribution, or more localized to one or both tails, or the center. Three pairwise comparisons were made (SAL - POL E, SAL - POL L, POL L - POL E) on the distributions for the duration for each call for each animal in the 5 minute recording period, thresholded between 5 ms and 300 ms (a range previously used to filter out noise(32)). A percentile bootstrap technique was used to derive confidence intervals based on differences in distribution at each decile of the distribution. This was then repeated to assess sex differences: the same comparisons were made in only males, and only females, followed by the same percentile bootstrap procedure. Sex differences were also investigated in the same way with the same pairwise comparisons as above for males and females separately.

#### 2.5.3. Partial least squares analysis

A partial least squares (PLS) analysis, previously applied by our group (11), was used to investigate putative brain-behaviour relationships between neonate neuroanatomy and USV behaviour. This is a multivariate technique for relating two sets of variables to each other by finding the optimal weighted linear combinations of variables that maximally covary with each other (33–35). The two variables used in this study were voxel-wise brain volumes (brain matrix) and USV call duration binned by decile of distribution (based on length), as well as sex, and number of pups per litter (described in **Table 1**)(behaviour/demographics matrix). The behaviour matrix was z-scored and correlated to the brain matrix to create a brain-behaviour covariance matrix. A singular value decomposition was applied to generate a set of orthogonal latent variables (LVs), which describe linked patterns of covariation between the input brain and behaviour matrices. Permutation testing and bootstrap resampling were applied to assess LV significance and reliability (further details in **Supplement 1.6.3**).

## 3. Results

### 3.1 Subtle neuroanatomical differences in neonate brain anatomy

Linear mixed-effects analysis of voxel wise volume difference revealed extremely subtle differences between SAL and POL E offspring (t=4.47, <20%FDR) wherein POL E offspring had a larger volume of a subregion of the right lateral amygdalar nucleus, a larger cluster of voxels in the right ventral hippocampus, and a smaller cluster of voxels in the right entorhinal cortex (**Figure 2**).

**Figure 2.**
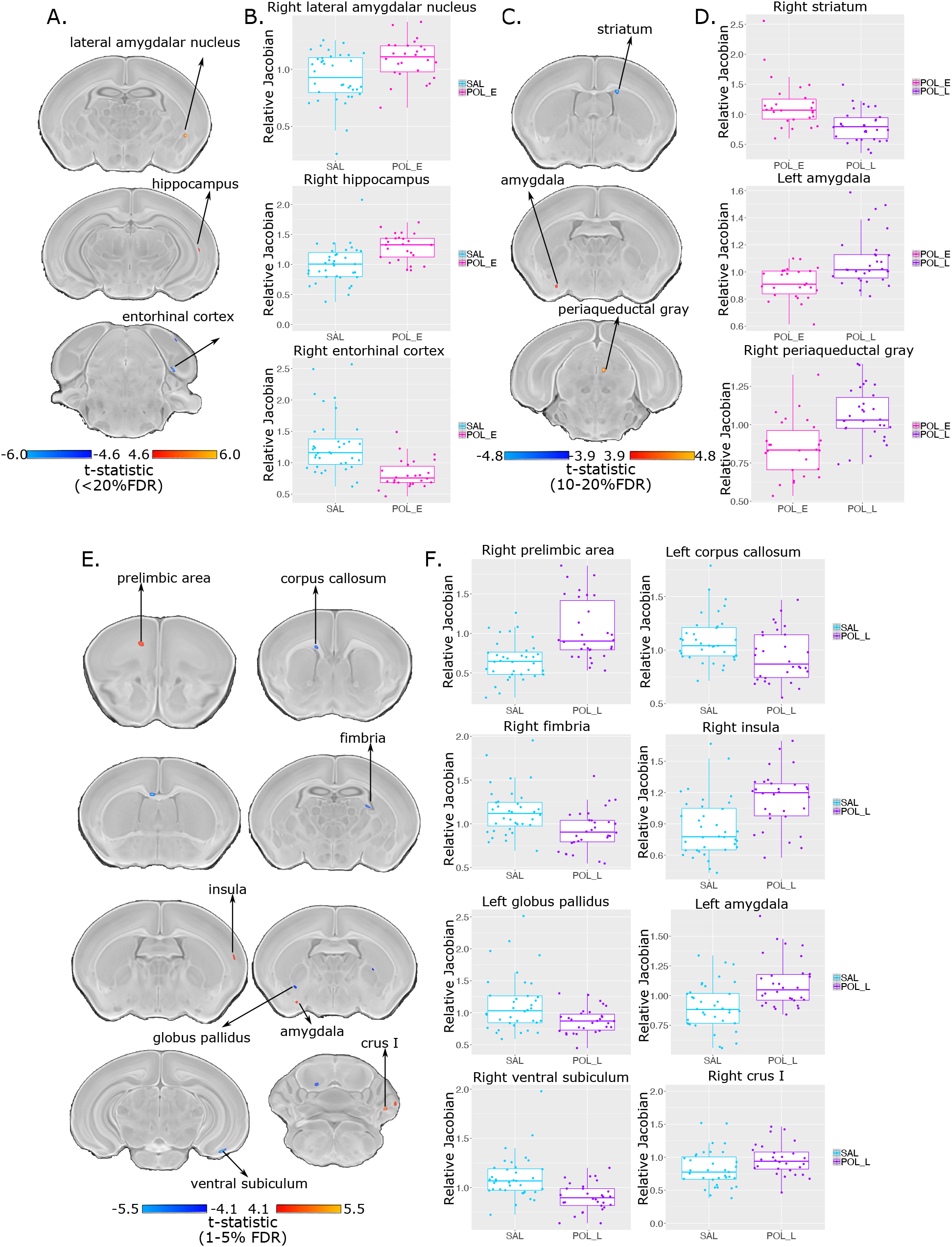
Neuroanatomical changes in the PND 8 neonate brain following GD 9 (**AB**) or GD 17 (**CD**) exposure. **A.** t-statistic map of group (POL E vs SAL) thresholded at 20% (bottom, t=4.60 to max, top, t=6.00) overlaid on the study average. **B.** Boxplot of peak voxels (voxels within a region of volume change showing largest effect) selected from regions of interest highlighted in **A**. **C.** t-statistic map of group (POL L vs POL E) thresholded at 20% (bottom, t=3.9) and 10% FDR (top, t=4.8) overlaid on the study average with peak voxels plotted in **D**. **E.** t-statistic map of group (POL L vs SAL) thresholded at 5% (bottom, t=4.10) and 1% FDR (top, t=5.50) overlaid on the study average with peak voxels plotted in **F**.

POL L offspring had more significant, yet focal changes relative to the SAL offspring (t=5.47, <1%FDR). Larger volume in a cluster within the right prelimbic area, left amygdala, right insula, and right Crus I of the cerebellum was observed in the POL L offspring. Conversely, volume decreases were observed within a subregion of the left corpus callosum and right fimbria, two white matter regions, as well as the right ventral subiculum and left globus pallidus in the POL L group (**Figure 2**). These results (brain maps) are shown at a less stringent threshold in **Supplement 2.2.1 and Supplementary figure 2).**

Finally, relative to the POL E group, the POL L group had smaller volume in subregions of the right striatum and larger volume in subregions of the left amygdala and right periaqueductal gray at a moderate significance threshold (t=4.80, <10%FDR) (**Figure 2**).

Post-hoc investigation of sex differences revealed subtle effects in the POL L group relative to SAL (t=4.23, <20%FDR); in the medial septum, pontine reticular nucleus, and cerebellum, volume for male POL L offspring was smaller than SAL, whereas the opposite was true for females, with larger volume for POL L than SAL offspring (**Supplement 2.2.2 and Supplementary figure 3)**. Additionally, the main effect of sex is represented in **Supplementary figure 4**(t=4.89, <1%FDR) in which we detect canonical sex differences in regions such as the medial preoptic area, the bed nucleus of the stria terminalis, and the medial amygdala (larger volume in males); these are similar to previously MRI-identified sex differences in the neonatal mouse brain (36). This suggests that we are sensitive to both profound and subtle changes, due to sex or MIA-exposure respectively.

### 3.2. Neonate ultrasonic vocalization behaviour results

No overall difference in mean call duration was observed in the ultrasonic vocalization data, however, data distribution and variance appeared different between groups. In order to better investigate potential differences in call duration, distributions for all calls made in the recording period, rather than mean call duration, were compared using the shift functions (**Figure 3A** and **Supplement 2.3**). The shift function revealed significant differences in distribution of call length between the SAL and POL E groups (**Figure 3B**). POL E offspring made significantly fewer calls at each decile of call length suggesting that overall they made shorter calls; more subtle differences were observed for deciles for shorter calls (decile 1, p=0.035, decile 2, p=0.003), and greater differences for longer duration calls (p<0.00001) (**Figure 3C**). The difference per decile is outlined in **Supplementary table 2**. Similar differences were observed between POL L and POL E offspring, again with POL E making significantly shorter calls identified by the shifted distribution, with the most significant differences observed for long calls (**Supplementary table 3**) (**Figure 3E**). Finally, there was no significant difference in distribution of call length between POL L and SAL offspring at any of the deciles (**Supplementary table 4**) (**Figure 3D**). Investigation of possible sex differences revealed that POL E females made significantly fewer long call across all deciles than SAL females, while there were no differences between POL E males and SAL males. Conversely, POL L females made significantly more calls than SAL females across most deciles of distribution, while no differences were observed in males (**Supplement 2.4** and **Supplementary figure 5**).

**Figure 3.**
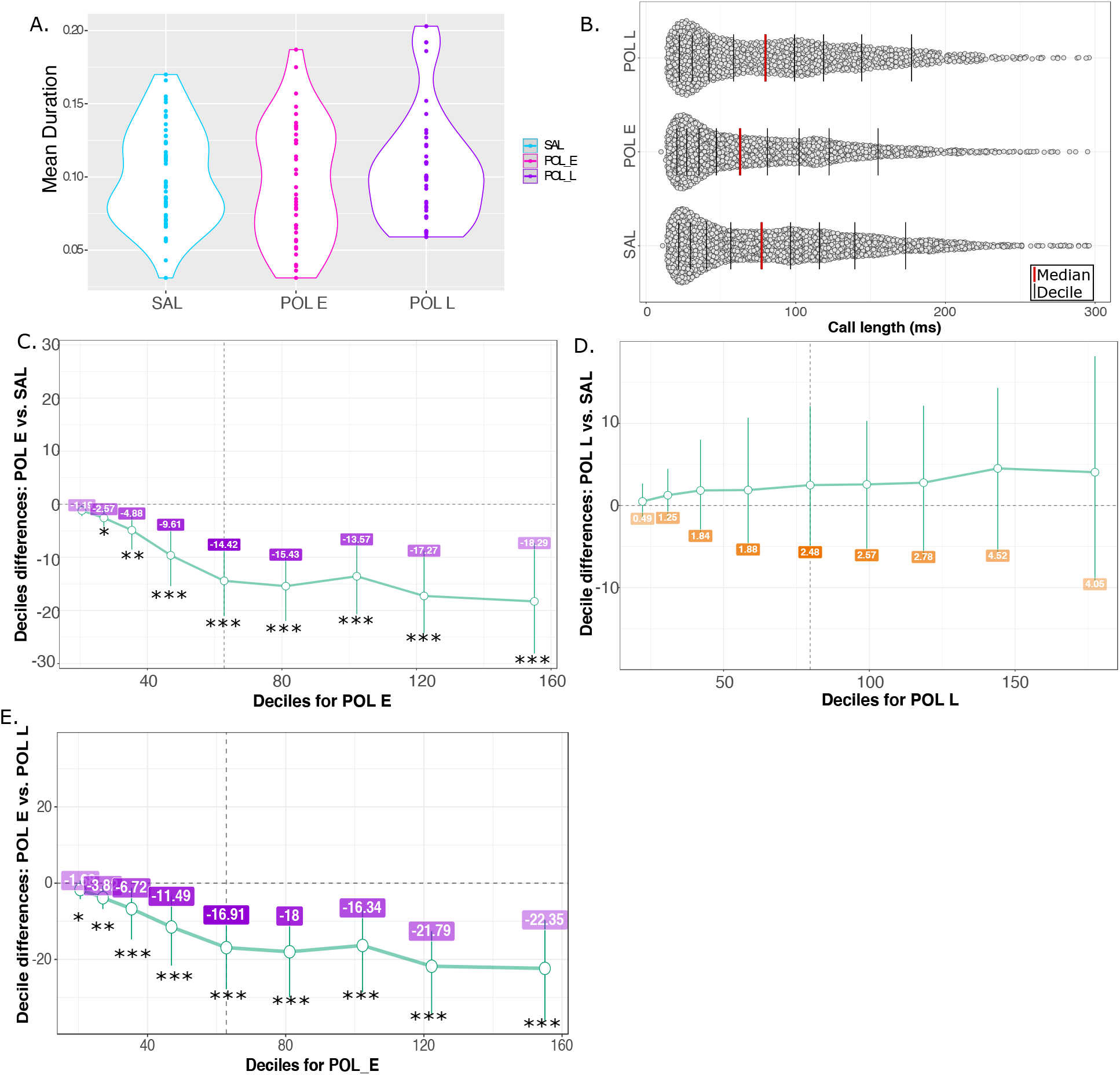
Results for ultrasonic vocalizations. **A.** Violin plot for mean call duration for each group (SAL, POL E, POL L) showing no overall differences in mean. **B.** Distribution of call length (ms) for all calls made by all mice per group in the 5-minute recording period. The red line identifies the median of the data, while each black bar denotes a decile of distribution. **C.** Percentile bootstrapping technique applied to identify the difference in decile between the POL E group and SAL, showing significantly fewer calls made by the POL E group across the range of distributions. **D.** Percentile bootstrapping analysis reveals no significant difference between distributions for POL L relative to SAL as error bars cross the zero line. **E.** Percentile bootstrapping analysis reveals significantly fewer calls for POL E group relative to POL L across all deciles. *p<0.05, **p<0.01, ***p<0.001

### 3.3. Neonate USV-brain PLS

PLS analysis of voxel-level volume changes and USV call frequency binned into deciles (as with the distribution analysis) yielded three significant latent variables (LV)s. The first accounted for 46% covariance between matrices (p=0.003) (**Figure 4A**). This showed a pattern of increased call frequency associated with larger volume in the left thalamus, left lateral septum, left auditory cortex, left ventral hippocampus, and fourth ventricle, and smaller volume in the right primary motor and somatosensory cortex, right amygdala, right and left corpus callosum, right and left thalamus (**Figure 4B,C**). The three groups load similarly onto the latent variable; however, the POL E and POL L groups seem to express the pattern slightly more than the SAL group (**Figure 4D**). The second LV accounted for 23% of the covariance (p=0.04) (**Figure 4E**) describing a pattern of increased corpus callosum, cingulate cortex, and subiculum volume, decreased thalamic and cerebellar volume associated with a greater number of shorter calls, and female sex (**Figure 4F,G**). Again, the POL E and POL L group load onto the latent variable slightly more than the SAL group (**Figure 4H**). Finally, the third LV is described in **Supplement 2.5** and **Supplementary figure 6** since it only accounted for 10% of the covariance (p=0.02) and the brain-behaviour covariation did not differentiate between groups.

**Figure 4.**
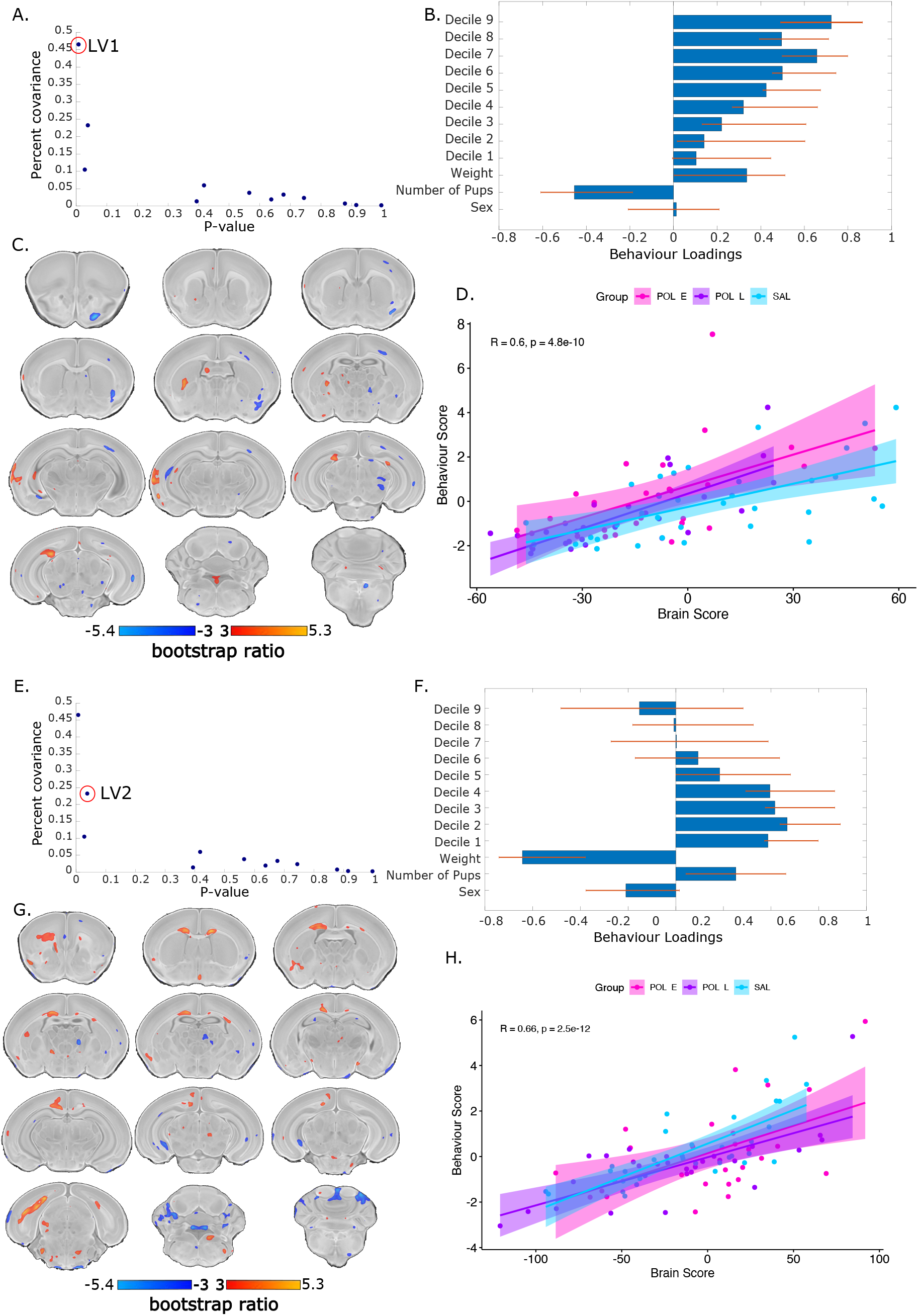
Partial least squares (PLS) analysis results for first and second significant latent variables (LV). **A**. Covariance explained (y-axis) and permutation p-values (x-axis) for all 12 LVs in the PLS analysis. LV1 is circled in red (p=0.003, %covariance=46%). **B.** Behaviour weight for each decile of distribution for the USV calls included in the analysis showing how much they contribute to the pattern of LV1. Singular value decomposition estimates the size of the bars whereas confidence intervals are estimated by bootstrapping. Bars with error bars that cross the 0 line should not be considered as significantly contributing to the LV. **C**. Brain loading bootstrap ratios for the LV1 deformation pattern overlaid on the population average, with positive bootstrap ratios in orange yellow (indicative or larger volume), and negative in blue (indicative of smaller volume). Colored voxels make significant contributions to LV1. **D**. Correlation of individual mouse brain and behaviour score, color coded by treatment group with a trend line per group. POL E offspring (magenta) express this pattern more strongly than SAL and POL L groups. **E.** LV2 is circled in red on the same plot as in **A** (p=0.04, %covariance=23%). Behaviour weights (**F**), brain weights (**G**), and brain-behaviour correlations (**H)** represented for LV2 as for LV1.

## 4. Discussion

Studies investigating the long-term impact of MIA-exposure on offspring development have identified a number of latent neuroanatomical, cellular, and behavioural abnormalities associated with this risk factor; many of these alterations overlap with those identified in neurodevelopmental disorders such as schizophrenia and autism spectrum disorder (37–39). While the majority of rodent MIA models have reported abnormalities in adolescent or adult offspring, here we investigate the less studied neonatal period. Based on the early life emergence of neurodevelopmental disorders such as autism spectrum disorder, as well as data from other preclinical studies in the rabbit and rhesus monkey (16,17), we hypothesized that neuroanatomical and behavioural deficits due to MIA-exposure would already be detectable in the neonatal period. Further, we hypothesized the timing of MIA-exposure would modulate the severity of offspring outcomes based on variation in neurodevelopment, and our previous findings (11). We leveraged high-resolution *ex vivo* MRI to characterize the effects of MIA-exposure either early or late in gestation on the neonatal brain anatomy in conjunction with an assay of communicative behaviours at PND 8.

Here, we observed subtle neuroanatomical changes due to MIA-exposure in the neonate at PND8. In our previous investigations of MIA-exposure on brain development, we observed MRI-detectable volume changes in the embryo brain due to both early and late exposure using a similar sample size. For early exposed embryos (GD9) volume reductions were observed in the hippocampus, globus pallidus, thalamus and cerebellum, while volume increases were observed in the prelimbic area, lateral septum, subventricular zone, and caudate putamen amongst others. In contrast, late exposed embryos (GD17) had striking brain-wide volume increases, particularly in the basal ganglia, hippocampus, cortex, corpus callosum, thalamus, and cerebellum (10). Furthermore, we previously identified significant deviation in neurodevelopmental trajectories of early MIA-exposed offspring in the adolescent and early adult period, again with a similar sample size, wherein accelerated volume increase, followed by a normalization were observed in the hippocampus, anterior cingulate cortex, striatum, and lateral septum, amongst other regions; in contrast, late-exposed offspring displayed only subtle deviations in trajectory (11). Thus, it is possible that the changes detected in the embryo brain are a result of acute remodeling in response to the increased inflammation *in utero*, but that these changes resolve in the neonatal period. The changes detected in the embryo period may be indicative of some latent disruptions that lead to aberrant neurodevelopment in later stages of adolescence and early adulthood. It is possible that the observations we made at PND8 capture some crossing trajectories of aberrant neurodevelopment, as depicted in **Figure 5**.

**Figure 5.**
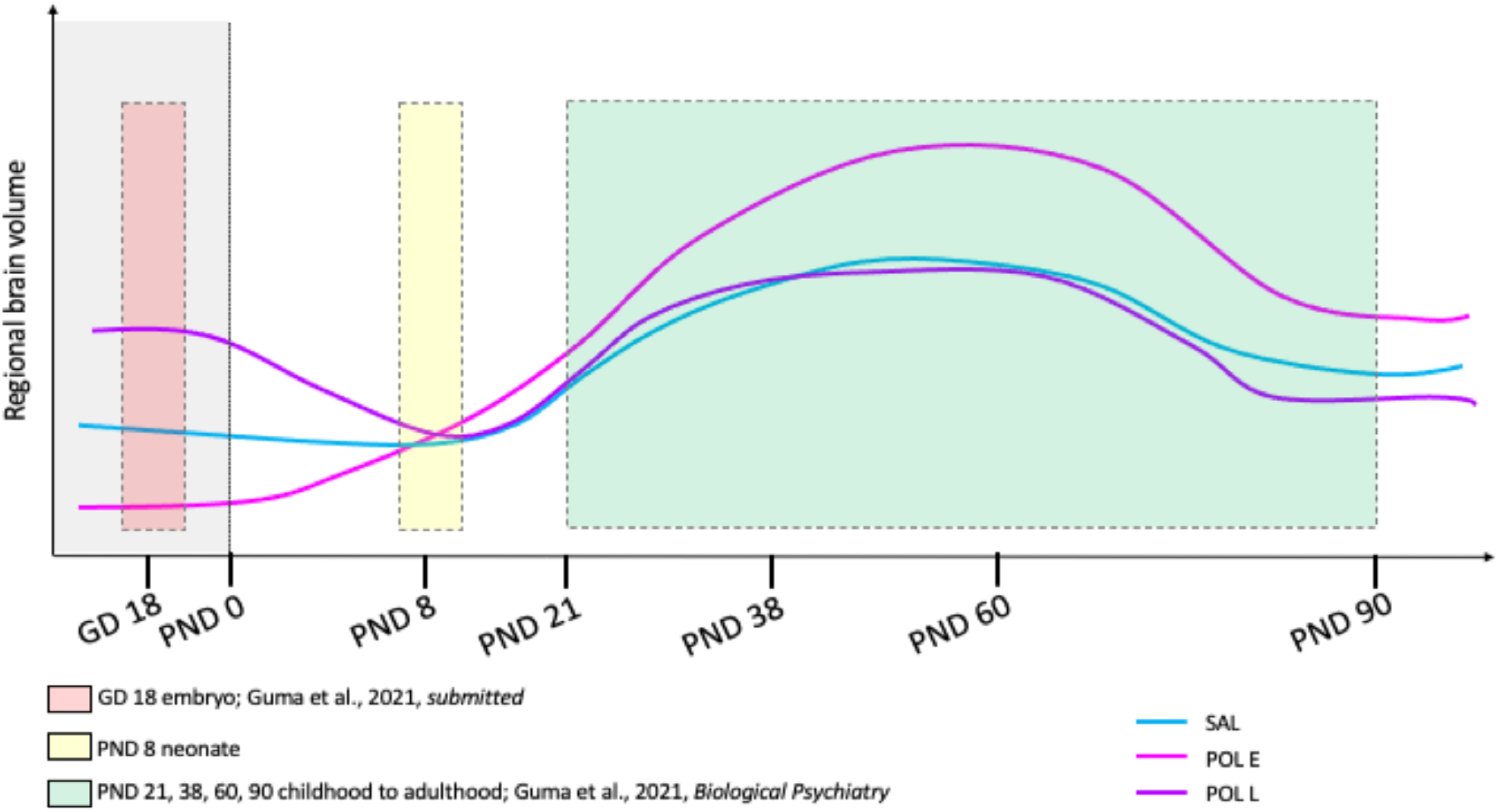
Hypothetical neurodevelopmental trajectories for the three examined groups, SAL, POL E, and POL L based on the work presented in this manuscript (PND8) as well as our previous investigation of GD18 embryo brain volume (10) and longitudinal brain development from PND21 to 90.

In contrast with our findings, brain imaging studies report neuroanatomical and functional changes associated with chronic exposure to inflammation *in utero* in human neonates (13–15). Furthermore, neonatal rhesus monkeys whose mothers were exposed to maternal immunoglobulin G (isolated from human mothers whose children had been diagnosed with ASD) in the first two trimesters displayed acceleration in brain growth, driven by white matter expansion particularly in the frontal and occipital lobes, and displayed behavioural features parallel to ASD (17). MIA-exposure in late gestation in the rabbit has also been shown to increase neuroinflammation in the first two postnatal weeks of life (measured by TSPO PET imaging) (16,40), and to decrease cortical serotonin binding (41). In addition to MIA-exposure, ASD may also be associated with alterations in neuroanatomy in the neonatal period, with differences detectable as early as 6 months of age (42). Differences in the nature of MIA-exposure may explain some of these differences, as both the human and rhesus monkey studies identified differences following more chronic inflammation. Differences in neurodevelopment between species could explain some of these discrepancies in findings (43). Naturally, the relatively minor alterations here require further investigation at both the structural and functional levels.

In an attempt to better understand how the timing of MIA-exposure may differentially impact neurodevelopment, it is important to consider the possible processes it may be interfering with. In the developing mouse brain, the neural tube forms, and neurogenesis begins at GD 9, corresponding with our early injection timing, and ends by GD15, prior to the late injection timepoint. Microglia also begin to colonize the brain at GD 9, making this developmental period sensitive to potential inflammatory insults such as those elicited by MIA (8,44,45). Thus, it is possible that MIA-exposure at GD9 may induce more lasting changes to cell proliferation and the development of the neuroinflammatory system. By GD 17, many neurodevelopmental processes are complete, however cortical cell migration and myelination are ongoing (46). In the GD17-exposed neonates we did observe subtle volume alterations to cortical regions, such as the prelimbic and insular cortices, as well as white matter regions such as the corpus callosum and fimbria. Finally, PND 8, the age at which we evaluated offspring brain anatomy and behaviour, likely corresponds to a human infant at term as myelination and synaptogenesis are underway during this window (47).

Social and communicative deficits are central to the pathology of many neurodevelopmental disorders, such as ASD. We experimentally modeled these impairments by measuring ultrasonic vocalizations in our MIA-exposed neonates, and found that early exposed groups made significantly fewer long calls which seem to be driven by the female offspring. Although the findings in the literature are fairly heterogeneous regarding the directionality of USV effects (i.e. increases or decreases due to MIA-exposure), previous groups have also observed decreases in neonatal vocalization in rodent MIA-models consistent with our findings (25), as well as other animal models for neurodevelopmental disorders(48,49). Interestingly, some studies have also investigated putative sex-dependent differences in this behaviour but observed greater alterations in male exposed offspring rather than in females as we observed(24,50).

Understanding the relationship between neuroanatomical abnormality and behaviour is critical and may be a useful avenue for understanding psychiatric disorders along dimensions rather than categories (51). We implement the multivariate technique, PLS, to investigate the relationship between voxel-level volume changes and USV call frequency. This allowed us to identify that neonates who made more frequent longer calls had a different brain anatomy than those that made more frequent shorter calls. Importantly, longer calls were associated with larger volume in the ventral hippocampus, lateral septum, auditory cortex, thalamus, and fourth ventricle and smaller volume in somatomotor cortices, corpus callosum, thalamus, and amygdala (based on the first latent variable with the highest covariance explained). In contrast, short calls were associated with larger volume in the corpus callosum, cingulate cortex, and subiculum, and smaller thalamic and cerebellar volume. Previous studies have found that neurotypical pup calls tend to be clustered in short sequences, thus, longer vocalizations may be indicative of some abnormal development or distress (52). Importantly, these patterns may be more expressed in both early and late MIA-exposed neonates. These results suggest that different brain regions may be involved in the production of short versus long vocalization. Further, some regions, such as the thalamus seem to be implicated across all patterns, suggesting that it may play a central role in communicative abilities.

The results presented in this manuscript should be considered in light of their limitations. The strategy we employed, leveraging high-resolution *ex vivo* MRI (rather than longitudinal) was selected as we thought it would be more sensitive at detecting neuroanatomical differences in the brain of neonate mice prenatally exposed to MIA. We expected to find more striking neuroanatomical alterations, but only recovered subtle, focal effects. We did observe more striking differences in neuroanatomy, particularly in the late-exposed offspring at a more lenient correction threshold, which may suggest that a larger sample size may be required to identify subtle differences in neuroanatomy at this developmental stage. Based on previous work by Lerch and colleagues (53) we were sufficiently powered to detect a 3% volume change with 10 subjects per group. However, it is possible that the within-subject variability in the neonatal timepoint may be higher than in adulthood given the rapid rate at which the brain is developing. The coefficient of variation (standard deviation/mean) for voxel-wise brain volume does indeed show a higher range in the neonatal sample compared to the embryo sample (10) and to the adolescent timepoint from our longitudinal sample (11) (see Table 2). It is conceivable that a single timepoint here, when the brain is growing so rapidly, might be noisier due to very subtle differences in maturation rates that for the purposes of this study add to the noise (described in **Supplement 1.6.1**).

**Table 2.** Coefficient of variation presented as a range across all voxel-wise brain data from the neonate data (PND8) presented here as well as two previous publications (10,11)

|  | **SAL** | **POL E** | **POL L** |
| --- | --- | --- | --- |
| **Embryo (GD18)** | 0 to 0.848 | 0 to 1.130 | 0 to 0.905 |
| **Neonate (PND8)** | 0 to 1.796 | 0 to 1.642 | 0 to 1.744 |
| **Adolescence (PND38)** | 0 to 0.520 | 0 to 0.448 | 0 to 0.468 |

In future work, employing longitudinal imaging on larger groups of offspring throughout the neonatal period may provide more sensitivity in detecting developmental changes rather than the cross-sectional approach employed here. This approach requires a significant amount of development, which our scanner was not equipped for at the time that this experiment was designed. This approach has been successful at uncovering sex differences throughout mouse brain development (36), and in previous work by our group examining offspring development from weaning to adulthood (11). Furthermore, we only assayed communicative abilities, as this is one of the few complex behaviours testable in neonate mice. However, a more comprehensive assay of social and repetitive/stereotypic behaviours (tested later in offspring development) may provide a more comprehensive examination of putative deficits relevant to ASD pathology (54). Finally, although the results indicated more subtle brain structure changes than we had expected, this observation advances our understanding of MIA-exposure on brain development in light of changes reported from the embryonic (10) and adolescent/early adult periods (55).

In conclusion, we comprehensively examined the effects of prenatal MIA-exposure, a known risk factor for neuropsychiatric disorders, at two gestational timepoints on neonatal brain anatomy and behaviour. Neuroanatomical differences were mostly resolved by the neonatal period, where we observed deficits in communication in the early exposed offspring, again, with greater effects in female offspring. These findings show that MIA-exposure induces subtle changes to neonatal development, which may require further investigation, but aid in our understanding of how this risk factor increases the likelihood of developing neuropsychiatric illnesses later in life.

## Supporting information

Supplementary material, results, figures and tables

## Acknowledgements

The authors are grateful to the laboratory of Dr. Lalit Srivastava for lending us the Ultrasonic Vocalizations equipment, particularly to Teresa Joseph. Additionally, we would like to thank Lourdes de Cossio Fernandez for sharing her protocol for performing the behavioural assays. We would like to thank Drs Bruno Giros and Salah El Mestikawy for lending us their centrifuge. Finally, the authors would like to acknowledge their funding bodies, including the Canadian Institute of Health Research and Healthy Brains for Healthy Lives for providing support for this research. Additionally, we would like to thank the Fonds de Recherche du Québec en Santé for providing salary support for EG and MMC, as well as the Kappa Kappa Gamma Foundation of Canada for supporting EG’s salary.

