## Supplementary material, results, figures and tables for "Subtle alterations in neonatal neurodevelopment following early or late exposure to prenatal maternal immune activation"

### Supplementary materials

#### 1. Supplementary methods

##### 1.1 Animals and maternal immune activation (MIA)

C57BL/6J mice were used throughout the study and were bred in our facility under a 12-hour light cycle (8am-8pm) with food and water access *ad libitum*. For timed mating procedure one male and one female of breeding age (8-12 weeks) were placed in a new cage in the afternoon, checked for appearance of plug, weighed, and separated the following morning. When a seminal plug was observed, this was considered gestational day (GD) 0 (mice were allowed only 1 night together to improve the accuracy of GD 0 detection).

For neonate sample collection, pregnant dams were randomly assigned to one of four treatment groups: (1) poly I:C (P1530-25MG polyinosinic:polycytidylic acid sodium salt TLR ligand tested; Sigma Aldrich) (5mg/kg, intraperitoneally) at gestational day (GD) 9 (POL E; n=6), (2) saline at GD9 (SAL E; n=4), (3) poly I:C at GD17 (POL L; n=6), (4) saline at GD17 (SAL L; n=4). GD9 corresponds roughly to the end of the first trimester in human gestation and GD17 corresponds to the end of the second trimester(Semple et al. 2013; Clancy, Darlington, and Finlay 2001). All injections were formed at 12pm (+/-1hour). All procedures were approved by McGill University's Animal Care Committee under the guidelines of the Canadian Council on Animal Care.

In a separate group of pregnant dams, poly I:C or saline was injected as described above (GD 9-POL: n=3 batch 1, n=4 batch 2; GD 17-POL n=3 batch 1, n=4 batch 2; GD 9-SAL n=5, GD 17-SAL n=3). Three hours following injection, dams were sacrificed by decapitation without euthanasia, and trunk blood was collected in a 1.5 mL Eppendorf tube. The blood was allowed to coagulate at room temperature for 30 minutes, and then centrifuged for 10 minutes (4 ℃, 2000 revolutions per minute). Serum was collected and stored at -80 ℃ until samples were shipped to the University of Maryland Core Cytokine Facility (http://www.cytokines.com/) for multiplex ELISA. Level of proinflammatory cytokines IL-6, TNF-α, IL-1β, and anti inflammatory cytokine IL-10 were measured to assess the immunostimulatory potential of our poly I:C.  We chose to use a separate group of dams to ensure we could collect enough serum for analysis, and so as not to introduce an additional stressful and potentially confounding experience for the dam. Detection ranges were as follows IL-6 (0.64-8000 pg/ml), TNF-α (0.64-3500 pg/ml), IL-1β (0.64-15000 pg/ml), IL-10 (0.64-20000 pg/ml).

##### 1.2 Brain sample preparation

On postnatal day (PND) 8, ~1 hour following behavioural testing neonates were given an i.p. injection of ketamine, xylazine, and acepromazine at a dose of 0.1 mL/100 g before being transcardially perfused. Perfusion was performed using 1x PBS solution with heparin, followed by 4% PFA with 2% Gadolinium in PBS. Skulls were extracted containing the brains of offspring, and post-fixed in the same PFA-gadolinium solution for 24 hours. After 24 hours, the brains were transferred from the PFA-PBS-gadolinium solution to a solution of 0.02% sodium azide, PBS, and 2% gadolinium.

##### 1.3 Magnetic resonance image acquisition

Prior to imaging, the samples were removed from the contrast agent solution, blotted and placed in 13-mm diameter plastic tubes filled with a proton-free susceptibility-matching fluid (Fluorinert FC-77, 3M Corp. St. Paul, MN). An anatomical scan was performed using a T2-weighted, 3D fast spin echo sequence using a cylindrical k-space acquisition (Nieman, et al., 2005) with TR/TE=350/12 ms, echo train length=6, two averages, field-of-view 20 mm x 20 mm x 25 mm, matrix size=504 x 504 x 630(Spencer Noakes, Henkelman, and Nieman 2017). Total imaging time was 14 hours producing images with 40 µm isotropic resolution. MR images of precision-machined phantoms were aligned towards a computed tomography (CT) scan of the same phantom to produce distortion correcting transformations to correct for geometric distortions in a coil-specific manner.

##### 1.4 MRI processing

An N4 correction for B1 bias field inhomogeneities(Tustison et al. 2010) was applied to brain images, and the background was set to zero using minc tools. Due to some perfusion artefacts, some brains presented with some extra matter that had similar intensity as the gray matter. To avoid misalignment during the registration, brain masks were manually segmented in order to target the automatic registrations to brain matter only (**Supplementary figure 1**).

Next, brain images of all neonates in the study were aligned using the antsMultivariateTemplateConstruction2.sh tool (see https://github.com/CoBrALab/twolevel_ants_dbm for implementation instructions)(Avants et al. 2011). Briefly, images were aligned using rigid registration (translation and rotation), affine (rigid, scaling, and shear), and finally, nonlinear registration to generate a precise anatomical alignment in an automated, minimally biased fashion. The output of this iterative registration procedure is a study-specific average against which groups can be compared, as well as deformation fields that map each individual subject to the average at the voxel level.

The Jacobian determinants of each deformation field provide a measure of volume difference at each voxel in the image relative to the average. Relative Jacobian determinants (used for statistical analysis in this work) explicitly model only the non-linear part of the deformations and remove residual global linear transformations (attributable to differences in total brain size). Prior to performing statistics, Jacobian determinants were blurred at 0.08 mm full-width-at-half-maximum to better conform to Gaussian assumptions for downstream statistical testing.

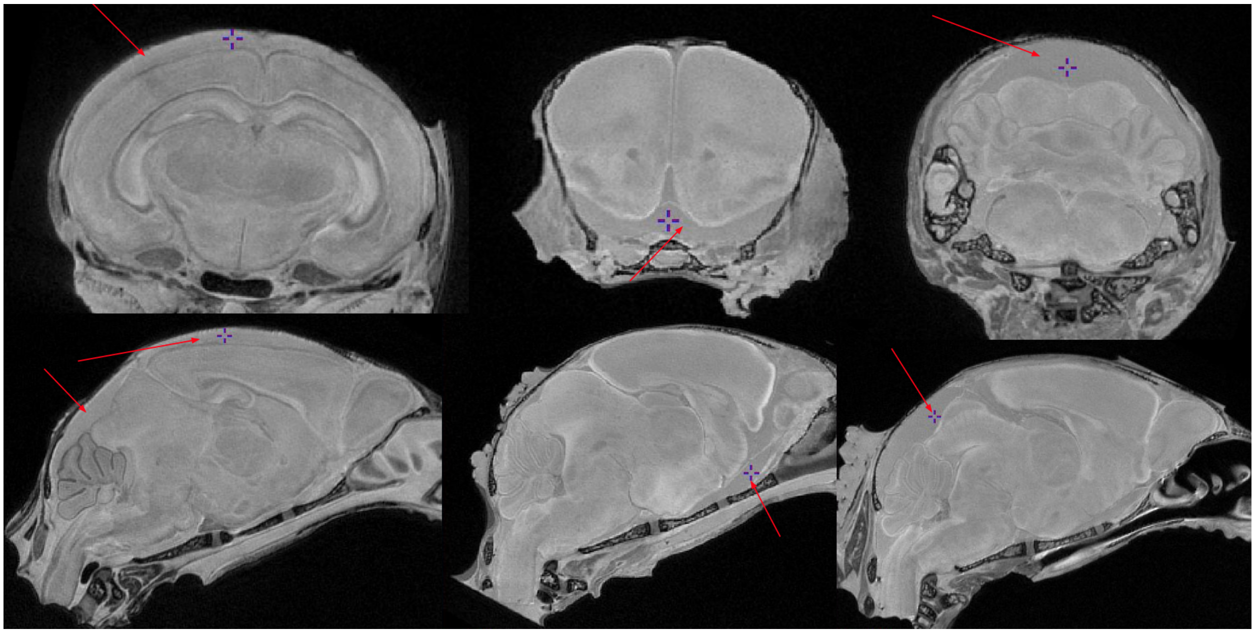

**Supplementary figure 1**. Examples of some of the perfusion artifacts observed in neonate scans. Whole-brain masks were manually segmented to ensure they did not bias registrations.

##### 1.5 Behavioural testing: ultrasonic vocalization task

Isolation-induced ultrasonic vocalizations of GD 9- and 17- poly I:C and saline exposed offspring were assessed with standard procedures(Cossío et al. 2017; Baharnoori, Bhardwaj, and Srivastava 2010). Testing was performed on PND 8, as the rate of calling peaks around this time in mouse pups (Scattoni et al., 2008). Fifteen minutes after the dam was removed from the home cage, each pup was individually transferred from the home cage to a plexiglass box (20.3 $cm^3$), where ultrasonic vocalizations were recorded continuously for 5 minutes using the Noldus UltraVox™ system (Noldus Information Technology, Leesburg, VA). An ultrasound detector (UltraSound Advice's Mini-3 bat detector) was tuned to detect incoming calls from 42 to 58 Hz. This was placed 15 cm above the bottom of the plexiglass box during each recording. A one-channel audio filter, which transmits a signal from the ultrasound detector to the computer if the signal is within the defined frequency amplitude range was set to detect calls if they lasted at least 5 ms and breaks in calls that lasted at least 1 ms.

This allowed us to maximize sensitivity to sound increases but avoid background noise. After each completed acquisition, pups were removed from the acquisition chamber, weighed, and returned to their home cage with their littermates. The acquisition box was cleaned with 20% ethanol between each session. The duration of each individual call per session per animal was recorded (raw data), as were the following summary measures per animal: total number of calls, maximum and minimum duration of calls, maximum and minimum call interval. POL E = 23 males, 17 females, SAL E = 12 males, 10 females, POL L = 25 males, 14 females, SAL L = 14 males, 16 females.

#### 1.6 Statistical analyses

##### 1.6.1 Neuroimaging analysis

Our first goal was to determine whether there were neuroanatomical differences in our two control groups, SAL E (GD9) and SAL L (GD17). We ran a whole-brain (delimited by a brain mask) voxel-wise linear mixed effects model on the relative Jacobian determinant files of each subject with injection timing and sex as fixed effects, and litter and number of pups per litter as random intercept to account for litter-specific variation. A False Discovery Rate (FDR) correction was applied. The same procedure was performed to assess whether there were any differences between the control neonate offspring (batch was not a factor for this analysis). In both cases, there were no differences, and SAL E and SAL L offspring were combined and set as the reference group. Finally, sex differences were explored, investigating the interaction between group and sex, with the same covariates as above. The statistical models applied to the relative Jacobian determinants to assess for group differences, described in **5.8.7.1** are detailed below:

**Main model:** $Y_{subject,j} = \beta_0 + \beta_1 sexF_{subject,j} + \beta_2 groupPOL\_E_{subject,j} + \beta_3 groupPOL\_L_{subject,j} + \mathbf{b_1} litter\ size + \epsilon_{subject,j}$

**Sex interaction model: model:** $Y_{subject,j} = \beta_0 + \beta_1 sexF_{subject,j} + \beta_2 groupPOL\_E_{subject,j} + \beta_3 groupPOL\_L_{subject,j} + \beta_4 sexF{:}groupPOL\_E_{subject,j} + \beta_5 sexF{:}groupPOL\_L_{subject,j} + \ + \mathbf{b_1} litter\ size + \epsilon_{subject,j}$

Y= outcome measures (i.e. blurred absolute Jacobian determinants); $\beta_i$= fixed effect coefficient; $\beta_0$ = equation intercept; **b** = random predictor; **ϵ** = random error; **j** = repeated measure per subject; **:** = interaction; POL_E = early polyI:C group relative to SAL as reference; POL_L = late polyI:C group relative to SAL as reference; SexF = female sex relative to male as reference

Additionally, the coefficient of variation (standard deviation/mean) was calculated on a voxel-by-voxel level within the brain mask for the neonate sample, and our previously published samples (Elisa Guma et al. 2021; E. Guma, do Couto Bordignon, and Devenyi 2021) using the minc Summary() functions to calculate the mean and standard deviation and the minccalc tool to calculate the difference maps.

##### 1.6.2 Neonate USV data analysis

For the USV data, summary measures were first tested for normality using a Shapiro-Wilks test. Since the data were not normally distributed, means were first compared using the Kruskal Wallis test. We determined that this was not the most accurate way to represent our data. Therefore, instead of using the summary measures provided by Ultravox, we examined the duration for each individual call made by each animal in the 5-minute recording period.

##### 1.6.3 Partial least squares analysis

Input imaging and behavioural data were organized into two matrices X (imaging), and Y (behaviour), with subjects as rows and variables in columns. The behavioural matrix was first z-scored (mean subtracted from each column and divided by standard deviation). A covariance matrix was then computed from the brain and z-scored behavioural matrices to represent all voxel deformation values and behavioural measures per subject. Singular value decomposition (SVD) was then applied to the covariance matrix(Eckart and Young 1936). This yields a set of orthogonal latent variables (LVs), which are patterns that describe the relationship between the brain and behaviour data. From these you get a set of ‘brain scores’ describing how each brain variable weights into a given LV, a set of ‘behaviour scores’ describing how each behaviour variable weights into a given LV, and a singular value, which describes the proportion of variance explained by the LV.

***Permutation testing*** was applied to assess the statistical significance of each LV wherein the rows (subjects) of the brain data matrix were randomly shuffled (n=1000 repetitions) to generate a null distribution of possible brain-behaviour correlations. The random shuffling allows for dependencies between brain and behaviour to be nullified. SVD was then applied to these “null” correlations, generating a distribution of singular values under the null hypothesis. A p-value can then be generated by looking at the probability that a permuted singular value exceeds the original, non-permuted singular value(Zeighami et al. 2019; Patel et al. 2020).

Next, ***bootstrap resampling*** was applied to assess the contribution of individual variables (both voxel and behavioural metrics) to each LV. Subjects (rows for both brain and behaviour matrices) were randomly sampled and replaced (n=1000) to generate a set of resampled correlation matrices to which SVD was applied to generate a sampling distribution for each weight of the singular vectors. The ratio of each singular vector weight and its bootstrap-estimated standard error were used to calculate a “bootstrap ratio” for each voxel. Voxels that make large contributions to certain patterns can therefore be identified by large bootstrap ratios. For this analysis, bootstrap ratios were thresholded at values corresponding to 95% confidence interval, as previously done(Zeighami et al. 2019). Next, weighted ‘brain’ and ‘behaviour scores’ were projected onto individual patient data to generate patient-specific scores.

#### 2. Supplementary results

##### 2.1 Poly I:C injection does increase pro-inflammatory cytokines

We observed an increase in levels of pro-inflammatory cytokines IL-6, IL-1β**,** and IL-10, but not TNF-α, in a separate cohort of pregnant dams 3-hours post poly I:C injection with our first batch on GD 9 relative to saline control on GD 9. For the second batch of poly I:C injected on GD 9, all 4 pro-inflammatory cytokines were increased (IL-6, IL-1β, TNF-α) 3 hours post-injection, but the anti-inflammatory cytokine IL-10 was not.

Exposure to our first batch of poly I:C on GD 17 increased levels of pro-inflammatory cytokines IL-6 relative to all but one SAL L dam who had exceptionally high IL-6 values. No differences were observed for TNF-α, IL-1β, or IL-10 relative to GD 17 saline controls. Our second batch of poly I:C similarly only increased IL-6, and had no effect on the other cytokine levels, however, for IL-1β and IL-10 values were below the detection threshold (<0.64) (**Supplementary Table 5.S1)**.

**Supplementary Table 1.** Maternal serum cytokine levels for our 4 treatment groups, mean [range]; * below detection range

| | IL-1β (pg/ml) | IL-6 (pg/ml) | IL-10 (pg/ml) | TNF-α (pg/ml) |
|---|---|---|---|---|
| **SAL E (n=5)** | 21.58 [0.64-96.62] | 45.54 [30.58-56.59] | 21.86 [0.64-56.22] | 14.30 [0.64-26.05] |
| **SAL L (n=3)** | 4.62 [0.64-6.88] | 746.33 [22.2-2192.47] | 42.13 [8.30-83.37] | 17.76 [5.99-39.59] |
| **POL E batch 1 (n=3)** | 11.56 [9.80-12.95] | 3578.84 [3624.87-3740.81] | 44.05 [36.93-49.70] | 8.20 [7.91-8.35] |
| **POL L batch 1 (n=3)** | 5.45 [5.00-6.34] | 1298.69 [54.36-3173.25] | 16.73 [13.67-19.58] | 20.35 [14.21-31.18] |
| **POL E batch 2 (n=4)** | 16.26 [0.64-25.4] | 14086.22 [1000.00- 18033.31] | 76.79 [35.18-174.44] | 69.05 [7.02-131.31] |
| **POL L batch 2 (n=4)** | 0.64* [0.64-0.64] | 1503.49 [0.64-1000] | 0.64 [0.64-0.64] | 4.95 [0.64-14.98] |

#### 2.2 Additional MRI results for POL L Offspring

##### 2.2.1 Differences in neuroanatomy due the GD17-MIA exposure at a lenient FDR threshold

Given that the neuroanatomical alterations in the POL L group were significant but very focal, they were explored at a more lenient FDR threshold, between 5-20%, to get a better understanding of them (**Supplementary Figure 2)**. These are consistent with the results presented in **Figure 2,** but cover larger patches of neuroanatomy.

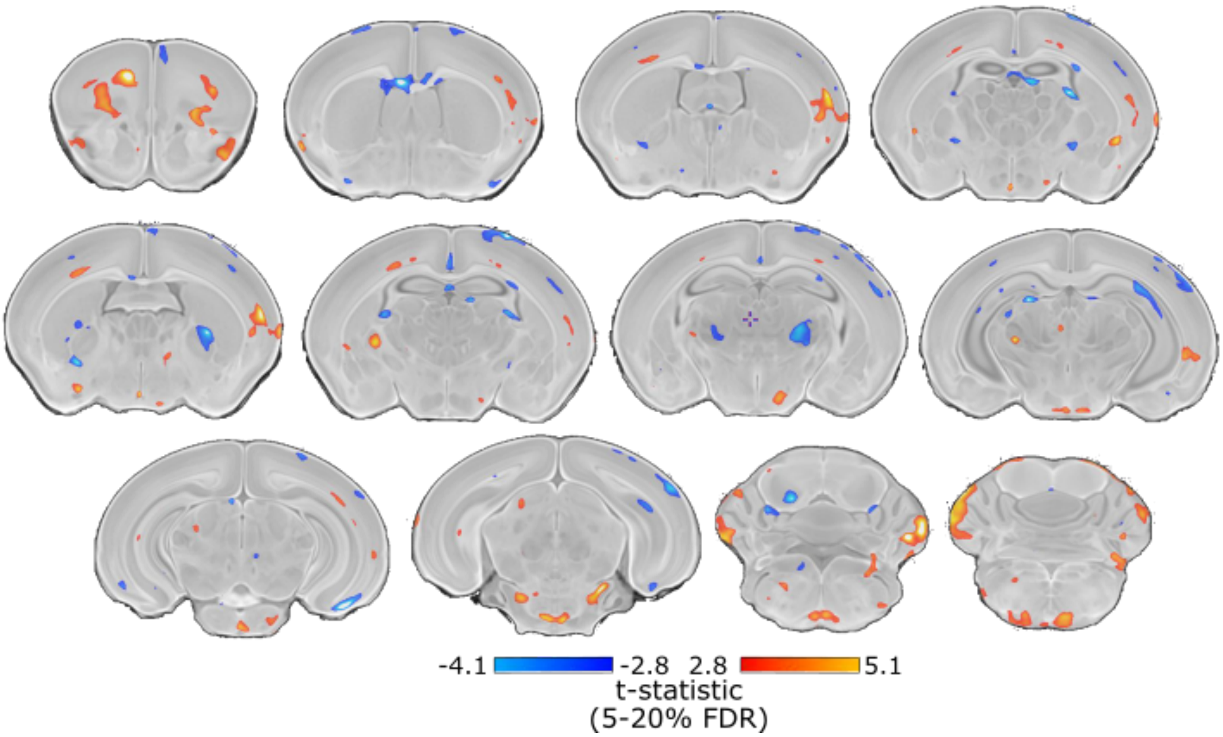


**Supplementary figure 2**. Neuroanatomical differences in the PND8 neonate brain following GD17 exposure at a more lenient threshold. t-statistic map of group (POL L vs SAL) thresholded between 5% and 20% FDR (top, t=-4.1, and bottom, t=2.8) overlaid on the study average.

##### 2.2.2 Sex differences neonate neuroanatomy

As described in the main text, post-hoc investigation of sex differences revealed subtle effects in the POL L group relative to SAL (t=4.23, q=0.20) wherein POL L males and smaller volume in the medial septum, reticular pontine nucleus, and cerebellum relative to SAL, with the opposite patter in females (**Supplementary figure 3)**. Additionally, as mentioned in the main text, the main effect of sex is plotted in **Supplementary figure 4** in which canonical sex differences are identified.

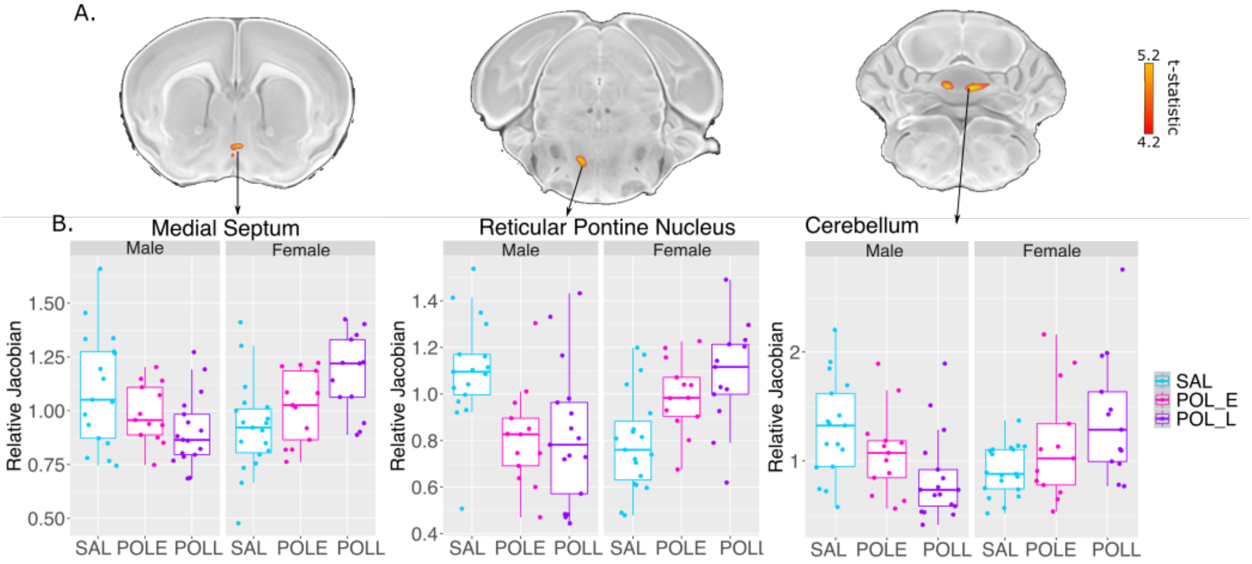


**Supplementary figure 3**. Sex differences in neuroanatomy in the PND8 neonate brain following GD17 exposure. **A.** t-statistic map of group (POL E vs SAL) thresholded at 20% (bottom, t=4.20) overlaid on the study average. **B.** Boxplot of peak voxels (voxels within a region of volume change showing largest effect) selected from regions of interest.

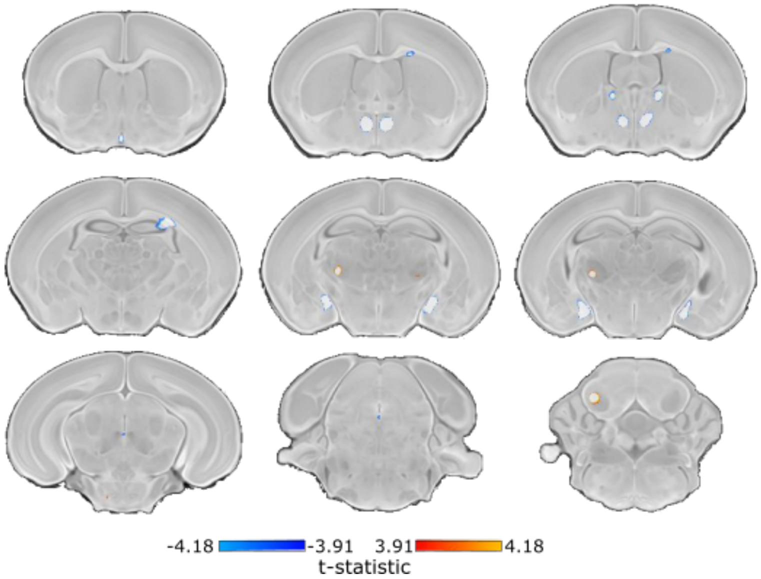


**Supplementary figure 4**. Sex differences in neuroanatomy in the PND8 neonate brain. **A.** t-statistic map of sex thresholded from 5% (top, t=4.18) to 10% FDR (bottom, t=3.91) overlaid on the study average. Regions in blue are larger in males and regions in yellow are larger in females.

##### 2.3 Analysis of summary USV data

Summary measures for USV data was not normally distributed based on the Shapiro-Wilks test for the following measures of interest: number of calls (W=0.821, p=2.43e-11), mean call duration (W=0.975, p=0.0155), and maximum call duration (W=0.849, p=3.07e-10). The Kruskall-Wallis rank sum test to compare group differences revealed no significant differences between the SAL, POL E, and POL L groups for number of calls (KW chi-squared=0.600, df=2, p=0.896), mean call duration (KW chi-squared=0.443, df=2, p=0.801), and maximum call duration (KW chi-squared=0.067, df=2, p=0.967).

#### 2.4 Sex differences in USVs

Investigation of possible sex differences revealed that POL E females made significantly fewer call across all deciles (all deciles, p<0.00001). POL E males made significantly more calls than SAL offspring at the first decile of distribution (p=0.025), but no significant differences were observed otherwise. POL L females made significantly more calls than SAL females from the second to eighth deciles (p>0.02), whereas POL L males, similar to the POL E males, made significantly more calls the SAL males only at the first decile of distribution (p=0.043) (**Supplementary figure 5; Supplementary tables 5-10**).

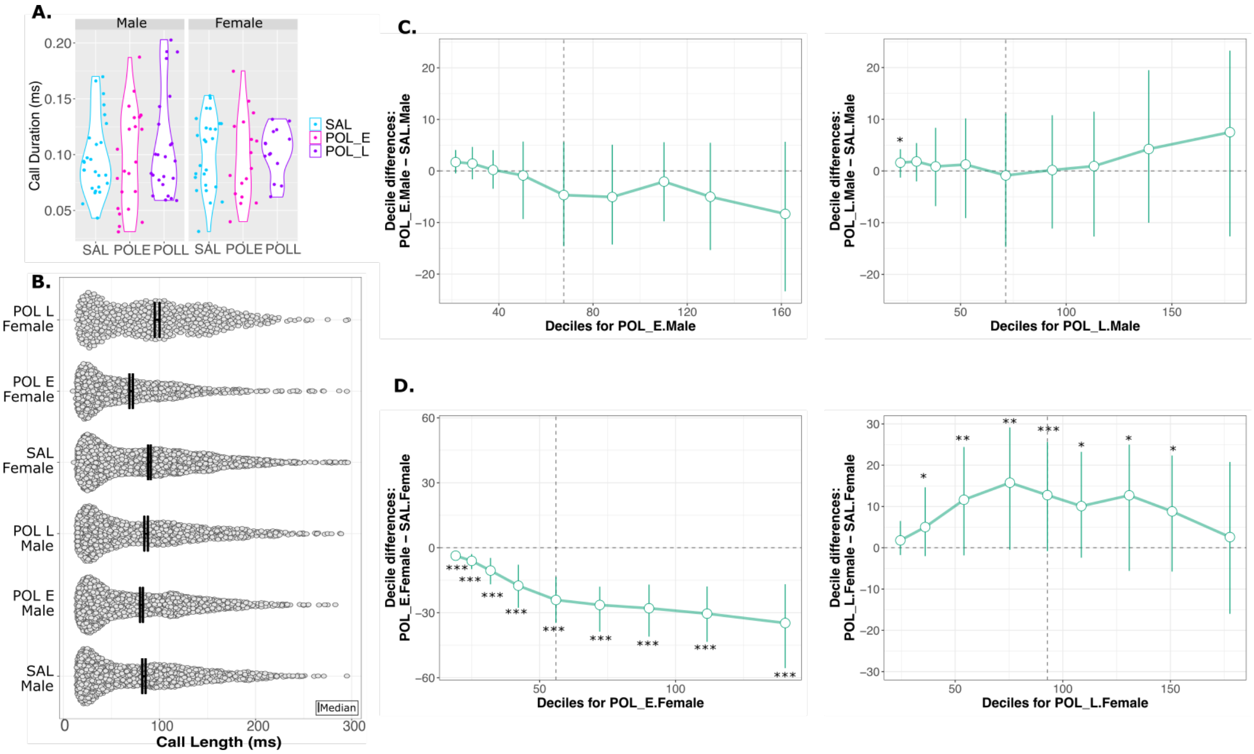


**Supplementary figure 5**. Sex differences in ultrasonic vocalizations. **A.** Violin plot for mean call duration for each group (SAL, POL E, POL L) split by males and females showing no significant differences. **B.** Distribution of call length (ms) for all calls made by all mice per group and per sex in the 5-minute recording period. The dark line identifies the median of the data, wherein the POL E females have a lower median than the other groups. **C.** Percentile bootstrapping technique applied to identify the difference in decile between the POL E and SAL males (right), and POL L and SAL males (right) showing no significant differences **D.** Percentile bootstrapping technique applied to identify the difference in decile between the POL E and SAL females (right), and POL L and SAL females (right) showing POL E females make significantly fewer calls than SAL females, with no differences in the POL L group. *p<0.05, **p<0.01, ***p<0.001

##### 2.5 Partial least squares results

The third significant latent variable described a pattern of increased striatal, thalamic, ventral hippocampal, septal, and amygdala volume and decreased dorsal hippocampus and corpus callosum volume with a greater number of short calls, and heavier body weight (**Supplementary figure 6)**.

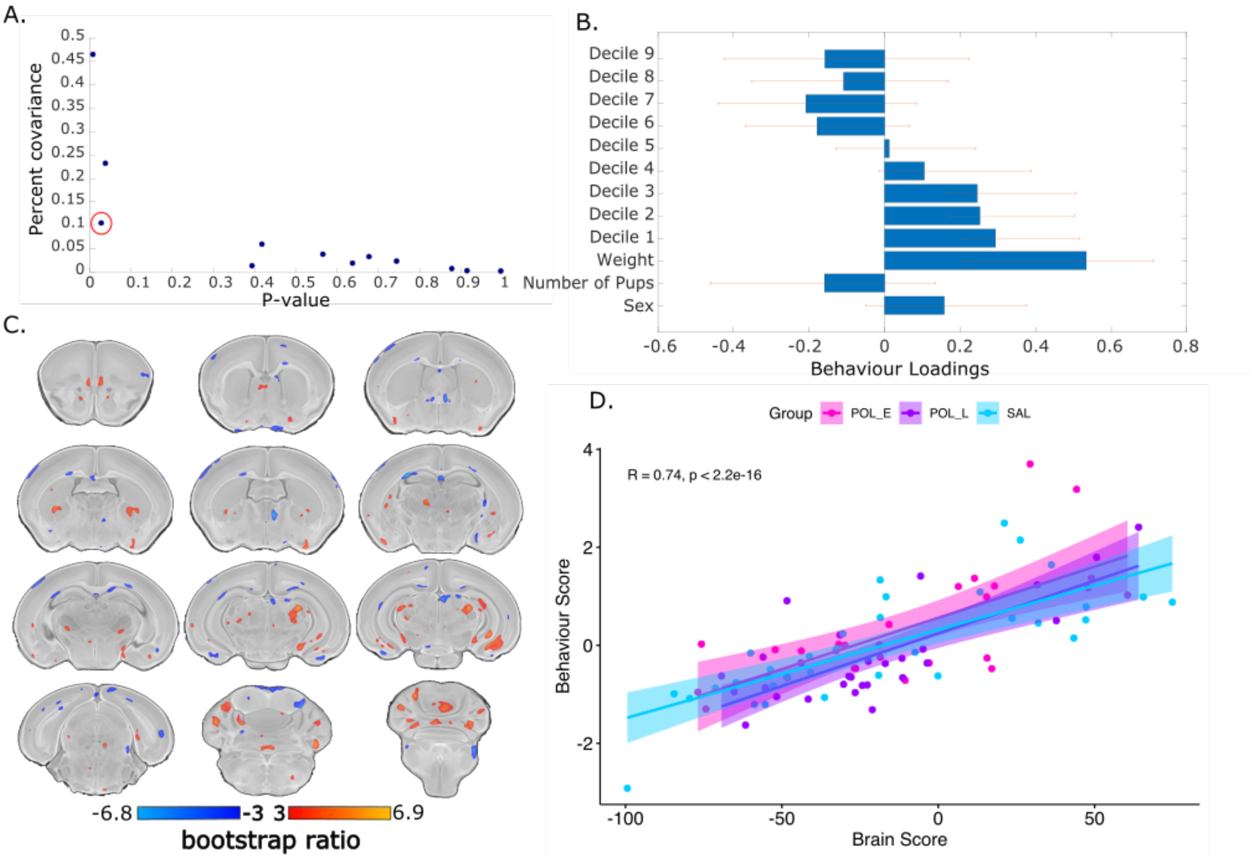


**Supplementary figure 6**. Partial least squares (PLS) analysis results for the third significant latent variable (LV3). **A**. Covariance explained (y-axis) and permutation p-values (x-axis) for all 12 LVs in the PLS analysis. LV3 is circled in red (p=0.02, %covariance=10%). **B.** Behaviour weight for each behavioural measure included in the analysis showing how much they contribute to the pattern of LV3. Singular value decomposition estimates the size of the bars whereas confidence intervals are estimated by bootstrapping. Bars with error bars that cross the 0 line should not be considered. **C**. Brain loading bootstrap ratios for the LV3 deformation pattern overlaid on the population average, with positive bootstrap ratios in orange-yellow (indicative or larger volume), and negative in blue (indicative of smaller volume). Colored voxels make significant contributions to LV3. **D**. Correlation of individual mouse brain and behaviour score, color coded by treatment group with a trend line per group. Early poly I:C (POL-E) offspring (magenta) express this pattern more strongly than the saline controls (SAL) and late poly I:C (POL-L) groups.

##### 2.6 Supplementary tables

For supplementary tables **2-5.14**, CI (confidence interval), p crit (uncorrected p-value), p-value (bootstrap corrected p-value); SAL (saline); POL E (early poly I:C group); POL L (late poly I:C group).

**Supplementary Table 2.** USV decile differences between SAL vs POL E

| Decile | SAL | POL E | Difference | CI lower | CI upper | p crit | p value |
|---|---|---|---|---|---|---|---|
| **1** | 21.714 | 20.525 | 1.1895 | 0.092 | 2.329 | 0.05 | 0.035 |
| **2** | 29.633 | 27.064 | 2.568 | 0.693 | 4.203 | 0.025 | 0.003 |
| **3** | 40.247 | 35.368 | 4.879 | 1.728 | 8.188 | 0.017 | <0.0001 |
| **4** | 56.539 | 46.926 | 9.613 | 3.755 | 15.523 | 0.013 | <0.0001 |
| **5** | 77.229 | 62.806 | 14.423 | 8.001 | 21.050 | 0.010 | <0.0001 |
| **6** | 96.559 | 81.133 | 15.426 | 9.131 | 21.920 | 0.0083 | <0.0001 |
| **7** | 115.807 | 102.242 | 13.565 | 6.503 | 20.644 | 0.0071 | <0.0001 |
| **8** | 139.515 | 122.2434 | 17.271 | 9.606 | 24.053 | 0.0063 | <0.0001 |
| **9** | 173.329 | 155.035 | 18.295 | 6.887 | 27.941 | 0.0056 | <0.0001 |

**Supplementary Table 3.** USV decile differences between POL L vs POL E

| Decile | POL L | POL E | Difference | CI lower | CI upper | p crit | p value |
|---|---|---|---|---|---|---|---|
| **1** | 22.208 | 20.525 | 1.684 | -0.241 | 4.049 | 0.0011 | 0.005 |
| **2** | 30.885 | 27.064 | 3.821 | 0.552 | 7.152 | 0.00056 | <0.0001 |
| **3** | 42.085 | 35.368 | 6.717 | 1.101 | 13.187 | 0.00037 | <0.0001 |
| **4** | 58.419 | 46.926 | 11.492 | 3.300 | 19.717 | 0.00028 | <0.0001 |
| **5** | 79.713 | 62.806 | 16.907 | 7.340 | 27.907 | 0.0002 | <0.0001 |
| **6** | 99.130 | 81.133 | 17.997 | 9.423 | 28.860 | 0.00019 | <0.0001 |
| **7** | 118.582 | 102.242 | 16.341 | 4.725 | 25.848 | 0.00016 | <0.0001 |
| **8** | 144.034 | 122.244 | 21.791 | 11.372 | 32.380 | 0.000134 | <0.0001 |
| **9** | 177.383 | 155.035 | 22.349 | 6.880 | 37.106 | 0.00012 | <0.0001 |

**Supplementary Table 4.** USV decile differences between SAL vs POL L

| Decile | SAL | POL L | Difference | CI lower | CI upper | p crit | p value |
|---|---|---|---|---|---|---|---|
| **1** | 21.714 | 22.208 | -0.494 | -2.556 | 1.450 | 0.0019 | 0.383 |
| **2** | 29.633 | 30.885 | -1.252 | -4.541 | 1.429 | 0.00069 | 0.187 |
| **3** | 40.247 | 42.085 | -1.838 | -8.138 | 3.594 | 0.0014 | 0.360 |
| **4** | 56.539 | 58.419 | -1.880 | -10.332 | 5.278 | 0.0056 | 0.478 |
| **5** | 77.229 | 79.713 | -2.484 | -11.559 | 6.460 | 0.0028 | 0.384 |
| **6** | 96.559 | 99.130 | -2.570 | -11.180 | 5.096 | 0.0008 | 0.327 |
| **7** | 115.807 | 118.582 | -2.776 | -12.281 | 5.927 | 0.0009 | 0.380 |
| **8** | 139.515 | 144.034 | -4.519 | -17.299 | 5.480 | 0.00062 | 0.142 |
| **9** | 173.329 | 177.383 | -4.054 | -18.817 | 7.7823 | 0.0011 | 0.333 |

**Supplementary Table 5.** USV decile differences between SAL males vs POL E males

| Decile | SAL Male | POL E Male | Difference | CI lower | CI upper | p crit | p value |
|---|---|---|---|---|---|---|---|
| **1** | 19.933 | 21.650 | -1.717 | -4.018 | 8 | 0.0056 | 0.025 |
| **2** | 27.340 | 28.767 | -1.427 | -4.615 | 1.663 | 0.008 | 0.219 |
| **3** | 37.337 | 37.550 | -0.212 | -3.886 | 3.472 | 0.05 | 0.883 |
| **4** | 51.184 | 50.306 | 0.878 | -5.670 | 9.181 | 0.025 | 0.777 |
| **5** | 72.241 | 67.573 | 4.6691 | -5.560 | 13.618 | 0.01 | 0.234 |
| **6** | 93.214 | 88.170 | 5.045 | -5.152 | 14.264 | 0.007 | 0.176 |
| **7** | 112.213 | 110.150 | 2.063 | -5.824 | 9.6212 | 0.017 | 0.552 |
| **8** | 134.826 | 129.820 | 5.005 | -5.852 | 15.361 | 0.01 | 0.215 |
| **9** | 169.970 | 161.640 | 8.329 | -6.038 | 24.231 | 0.006 | 0.115 |

**Supplementary Table 6.** USV decile differences between SAL males vs POL L males

| Decile | SAL Male | POL_L Male | Difference | CI lower | CI upper | p crit | p value |
|---|---|---|---|---|---|---|---|
| **1** | 19.933 | 21.535 | -1.602 | -3.652 | 0.778 | 0.0007 | 0.043 |
| **2** | 27.340 | 29.170 | -1.830 | -5.608 | 1.975 | 0.001 | 0.165 |
| **3** | 37.337 | 38.211 | -0.873 | -8.430 | 6.296 | 0.001 | 0.703 |
| **4** | 51.184 | 52.423 | -1.238 | -10.188 | 9.537 | 0.001 | 0.756 |
| **5** | 72.241 | 71.3778 | 0.863 | -11.778 | 13.412 | 0.003 | 0.893 |
| **6** | 93.214 | 93.391 | -0.177 | -11.402 | 9.847 | 0.006 | 0.961 |
| **7** | 112.213 | 113.1518 | -0.939 | -12.256 | 9.914 | 0.002 | 0.809 |
| **8** | 134.826 | 139.070 | -4.244 | -19.628 | 9.285 | 0.001 | 0.362 |
| **9** | 169.969 | 177.468 | -7.499 | -22.819 | 12.207 | 0.001 | 0.259 |

**Supplementary Table 5.S7.** USV decile differences between SAL females vs POL E females

| Decile | SAL Female | POL E Female | Difference | CI lower | CI upper | p crit | p value |
|---|---|---|---|---|---|---|---|
| 1 | 22.76142 | 19.11462 | 3.646794 | 1.533147 | 5.318447 | 9.92E-05 | <0.0001 |
| 2 | 31.06479 | 25.0127 | 6.052092 | 3.102647 | 10.140771 | 4.96E-05 | <0.0001 |
| 3 | 42.45443 | 31.93909 | 10.515341 | 3.182823 | 16.82793 | 3.31E-05 | <0.0001 |
| 4 | 59.66499 | 42.16741 | 17.497579 | 6.717054 | 26.904838 | 2.48E-05 | <0.0001 |
| 5 | 80.10018 | 56.00561 | 24.094573 | 12.569376 | 33.780367 | 1.98E-05 | <0.0001 |
| 6 | 98.53035 | 72.12251 | 26.407841 | 13.788924 | 37.519 | 1.65E-05 | <0.0001 |
| 7 | 118.16617 | 90.23947 | 27.926706 | 17.378152 | 39.514686 | 1.42E-05 | <0.0001 |
| 8 | 141.98812 | 111.587 | 30.401119 | 18.310493 | 43.222528 | 1.24E-05 | <0.0001 |
| 9 | 175.07448 | 140.33144 | 34.743037 | 16.7773 | 49.852391 | 1.10E-05 | <0.0001 |

**Supplementary Table 8.** USV decile differences between SAL females vs POL L females

| Decile | SAL Female | POL L Female | Difference | CI lower | CI upper | p crit | p value |
|---|---|---|---|---|---|---|---|
| **1** | 22.761 | 24.552 | -1.791 | -6.782 | 1.777 | 5.51E-06 | 0.185 |
| **2** | 31.065 | 36.080 | -5.0147 | -15.719 | 1.006 | 2.20E-06 | 0.013 |
| **3** | 42.454 | 54.088 | -11.633 | -27.299 | 0.829 | 1.57E-06 | 0.005 |
| **4** | 59.665 | 75.432 | -15.767 | -29.412 | 2.100 | 1.22E-06 | 0.002 |
| **5** | 80.100 | 92.849 | -12.7492 | -28.968 | -1.298 | 1.38E-06 | <0.0001 |
| **6** | 98.530 | 108.621 | -10.090 | -24.425 | 1.36145 | 1.84E-06 | 0.011 |
| **7** | 118.166 | 130.870 | -12.704 | -27.805 | 3.778 | 2.76E-06 | 0.013 |
| **8** | 141.988 | 150.794 | -8.806 | -23.538 | 3.433 | 3.67E-06 | 0.029 |
| **9** | 175.074 | 177.658 | -2.584 | -24.780 | 15.394 | 1.10E-05 | 0.614 |

**Supplementary Table 9.** USV decile differences between POL L males vs POLE males

| Decile | POL L Female | POL_E Female | Difference | CI lower | CI upper | p crit | p value |
|---|---|---|---|---|---|---|---|
| **1** | 21.535 | 21.650 | -0.115 | -2.942 | 2.264 | 8.93E-04 | 0.858 |
| **2** | 29.170 | 28.767 | 0.403 | -5.483 | 4.725 | 4.46E-04 | 0.742 |
| **3** | 38.211 | 37.550 | 0.662 | -5.016 | 7.429 | 2.98E-04 | 0.725 |
| **4** | 52.423 | 50.306 | 2.117 | -7.071 | 11.555 | 2.23E-04 | 0.474 |
| **5** | 71.378 | 67.573 | 3.805 | -12.423 | 15.268 | 1.49E-04 | 0.373 |
| **6** | 93.391 | 88.170 | 5.221 | -6.892 | 17.637 | 1.28E-04 | 0.192 |
| **7** | 113.152 | 110.150 | 3.002 | -9.600 | 15.952 | 1.79E-04 | 0.400 |
| **8** | 139.069 | 129.820 | 9.249 | -5.530 | 23.945 | 1.12E-04 | 0.041 |
| **9** | 177.468 | 161.640 | 15.828 | -1.051 | 30.105 | 9.92E-05 | 0.001 |

**Supplementary Table 10.** USV decile differences between POL L females vs POL E females

| Decile | POL L Female | POL E Female | Difference | CI lower | CI upper | p crit | p value |
|---|---|---|---|---|---|---|---|
| **1** | 24.552 | 19.115 | 5.438 | 1.371 | 10.853 | 1.10E-05 | <0.0001 |
| **2** | 36.079 | 25.013 | 11.067 | 4.826 | 18.369 | 5.51E-06 | <0.0001 |
| **3** | 54.088 | 31.940 | 22.149 | 6.850 | 37.049 | 3.67E-06 | <0.0001 |
| **4** | 75.432 | 42.167 | 33.265 | 16.423 | 45.459 | 2.76E-06 | <0.0001 |
| **5** | 92.849 | 56.006 | 36.844 | 22.777 | 49.530 | 2.20E-06 | <0.0001 |
| **6** | 108.621 | 72.123 | 36.498 | 24.534 | 51.766 | 1.84E-06 | <0.0001 |
| **7** | 130.870 | 90.240 | 40.631 | 20.594 | 54.509 | 1.57E-06 | <0.0001 |
| **8** | 150.794 | 111.587 | 39.207 | 24.807 | 53.958 | 1.38E-06 | <0.0001 |
| **9** | 177.658 | 140.331 | 37.327 | 15.605 | 60.514 | 1.22E-06 | <0.0001 |

Patel, Raihaan, Christopher J. Steele, Anthony G. X. Chen, Sejal Patel, Gabriel A. Devenyi, Jürgen Germann, Christine L. Tardif, and M. Mallar Chakravarty. 2020. “Investigating Microstructural Variation in the Human Hippocampus Using Non-Negative Matrix Factorization.” *NeuroImage* 207 (February): 116348.

Semple, Bridgette D., Klas Blomgren, Kayleen Gimlin, Donna M. Ferriero, and Linda J. Noble-Haeusslein. 2013. “Brain Development in Rodents and Humans: Identifying Benchmarks of Maturation and Vulnerability to Injury across Species.” *Progress in Neurobiology* 106-107 (July): 1–16.

Spencer Noakes, T. Leigh, R. Mark Henkelman, and Brian J. Nieman. 2017. “Partitioning K-Space for Cylindrical Three-Dimensional Rapid Acquisition with Relaxation Enhancement Imaging in the Mouse Brain.” *NMR in Biomedicine* 30 (11): e3802.

Tustison, Nicholas J., Brian B. Avants, Philip A. Cook, and James C. Gee. 2010. “N4ITK: Improved N3 Bias Correction with Robust B-Spline Approximation.” *2010 IEEE International Symposium on Biomedical Imaging: From Nano to Macro*. https://doi.org/10.1109/isbi.2010.5490078.

Zeighami, Yashar, Seyed-Mohammad Fereshtehnejad, Mahsa Dadar, D. Louis Collins, Ronald B. Postuma, Bratislav Mišić, and Alain Dagher. 2019. “A Clinical-Anatomical Signature of Parkinson’s Disease Identified with Partial Least Squares and Magnetic Resonance Imaging.” *NeuroImage* 190 (April): 69–78.
